# Effects of long-term high dose aspartame on body mass, bone strength, femoral geometry, and microbiota composition in a young and aged cohort of male and female mice

**DOI:** 10.1101/2024.01.02.573970

**Authors:** Erika L. Cyphert, Chongshan Liu, Angie L. Morales, Jacob C. Nixon, Emily Blackford, Matthew Garcia, Nicolas Cevallos, Peter J. Turnbaugh, Ilana L. Brito, Sarah L. Booth, Christopher J. Hernandez

## Abstract

**Background:** Recent reassessment of the safety of aspartame has prompted increased evaluation of its effect on the health of a range of tissues. The gut microbiome is altered by oral aspartame. One prior study suggested that changes in the microbiome caused by aspartame could influence the strength of bone in young skeletally developing mice. Here we ask how aspartame influences bone in mice of different age and sex.

**Objective:** The objective of this study was to determine the effect of aspartame on the bone strength and gut microbiota of young and aged mice.

**Methods:** Male and female C57Bl/6J mice were untreated or treated with a high dose of aspartame in their drinking water from 1 month of age until 4 (young cohort; n = 80) or 22 months (aged cohort; n = 52).

**Results:** In aged males, mice treated with aspartame had greater body mass, whole bone strength, and femoral geometry relative to untreated. Specifically, in aged males, aspartame led to 9% increase in body mass (p < 0.001), 22% increase in whole bone strength (p = 0.006), and 17% increase in section modulus (p < 0.001) relative to untreated mice. Aged males and females receiving aspartame had a different microbiota than untreated mice and a decreased abundance of *Odoribacter*. No differences in body mass, whole bone strength, or femoral geometry were associated with aspartame dosing in young males or young or aged females.

**Conclusions:** Aspartame treated aged males had greater whole bone strength and the effect appeared to be explained by greater body mass. Aspartame treatment did not alter whole bone strength in young males or young or aged females despite the aspartame having a similar effect on the microbiota of both aged males and females.

## Introduction

Consumption of aspartame is common and increasing in some populations. In the US 41% of adults consume aspartame daily, while in adolescents consumption of aspartame has increased 200% over the past two decades (1,2). While sweeteners such as aspartame were initially marketed as a zero-calorie alternative to sucrose, aspartame consumption has been linked to changes in liver function (3), neurological function (4,5), and glucose tolerance and obesity (6,7). However, the effect of aspartame on organ function is not consistent across studies. The World Health Organization has increased scrutiny of aspartame and recently classified it as a possible carcinogen (8). Furthermore, non-nutritive sweeteners like aspartame can alter the composition of the microbiota (6,7) and thereby affect organs throughout the body.

The composition of the gut microbiota has been shown to influence both bone quantity (amount of bone) and quality (microarchitecture, mineral/matrix composition, collagen crosslinking, etc.) (9–12). In a prior study, an aspartame-based sweetener was used to ensure adequate intake of antibiotic-laced water by mice. A control group of mice receiving only the sweetener resulted in 40% greater whole bone strength relative to mice not receiving aspartame by altering the mechanical properties of the bone matrix (not just bone size/density) (9). This finding was exciting in that it suggested that an intervention could increase the mechanical performance of bone matrix, a contributor to age-related fracture in aged populations that has not previously been explored as a therapeutic target (13). However, this prior study was limited in that it was not designed to examine the effects of the aspartame-based sweetener, it only examined male mice, and it included only one time point (collected at 4 months of age after 3 months of aspartame dosing). As a result, it remains unclear how chronic aspartame treatment influenced whole bone strength and geometry, or if the effect was limited to one sex or age.

The goal of this work was to determine the effect of aspartame on the gut microbiota and bone strength. Specifically, we sought to determine: 1) the effect of aspartame on bone strength and geometry across age and sex, 2) the effect of aspartame on the composition of the gut microbiota, and 3) which microbial taxa were associated with changes in bone strength following aspartame dosing.

## Methods

### Mouse strains

Animal procedures were approved by the local Institutional Animal Care and Use Committee. Male and female C57Bl/6J mice were purchased from Jackson Laboratory (Bar Harbor, ME, US) and bred using trio breeding. Pups were weaned at three weeks of age and randomly housed by sex (n = 3-4/cage). Sterile cages contained ¼-inch corn cob bedding (The Andersons’ Lab Bedding, Maumee, OH, US) and cardboard hut (Ketchum Manufacturing, Brockville, Canada).

### Diet and aspartame sweetener

Mice received standard laboratory chow (Teklad LM-485 Mouse/Rat Sterilizable Diet, Envigo Diets, Madison, WI, US; ingredient list outlined in **Supplementary Table 1**) and reverse osmosis sterilized water *ad libitum* with or without 10 g/L zero-calorie sweetener (Equal: aspartame, dextrose, maltodextrin, acesulfame potassium; Merisant Company, Chicago, IL, US) (9). Aspartame sweetener laced water was freshly prepared and replenished every three days.

### Study design

The study included both male and female and both young (4 months of age) and aged (22 months of age) animals. Pups were divided into four groups based on age and treatment (number of animals in each treatment group is outlined in **Supplementary Table 2**): 1) chronic aspartame dosing from 1-4 months of age, 2) untreated control 4 months of age, 3) chronic aspartame dosing from 1-22 months of age, and 4) untreated control 22 months of age (**Figure 1A**). A sample size of 12 animals per group had a statistical power of 0.80 to detect an effect size of 0.88 with α = 0.05. Larger sample sizes were bred to control for age-related attrition, but all available animals were used for the study resulting in variance in group size. Dosing consisted of reverse osmosis water containing zero-calorie aspartame sweetener (10 g/L; freshly prepared every 3 days) (9). Weekly mixing of bedding between cages of the same sex and treatment group was applied from 1-3 months of age to reduce cage to cage variation in the microbiota (14). Mice were euthanized at either 4 or 22 months of age and femora were dissected and preserved by wrapping in gauze soaked with phosphate buffered saline and plastic wrap stored at −80°C.

**Figure 1:**
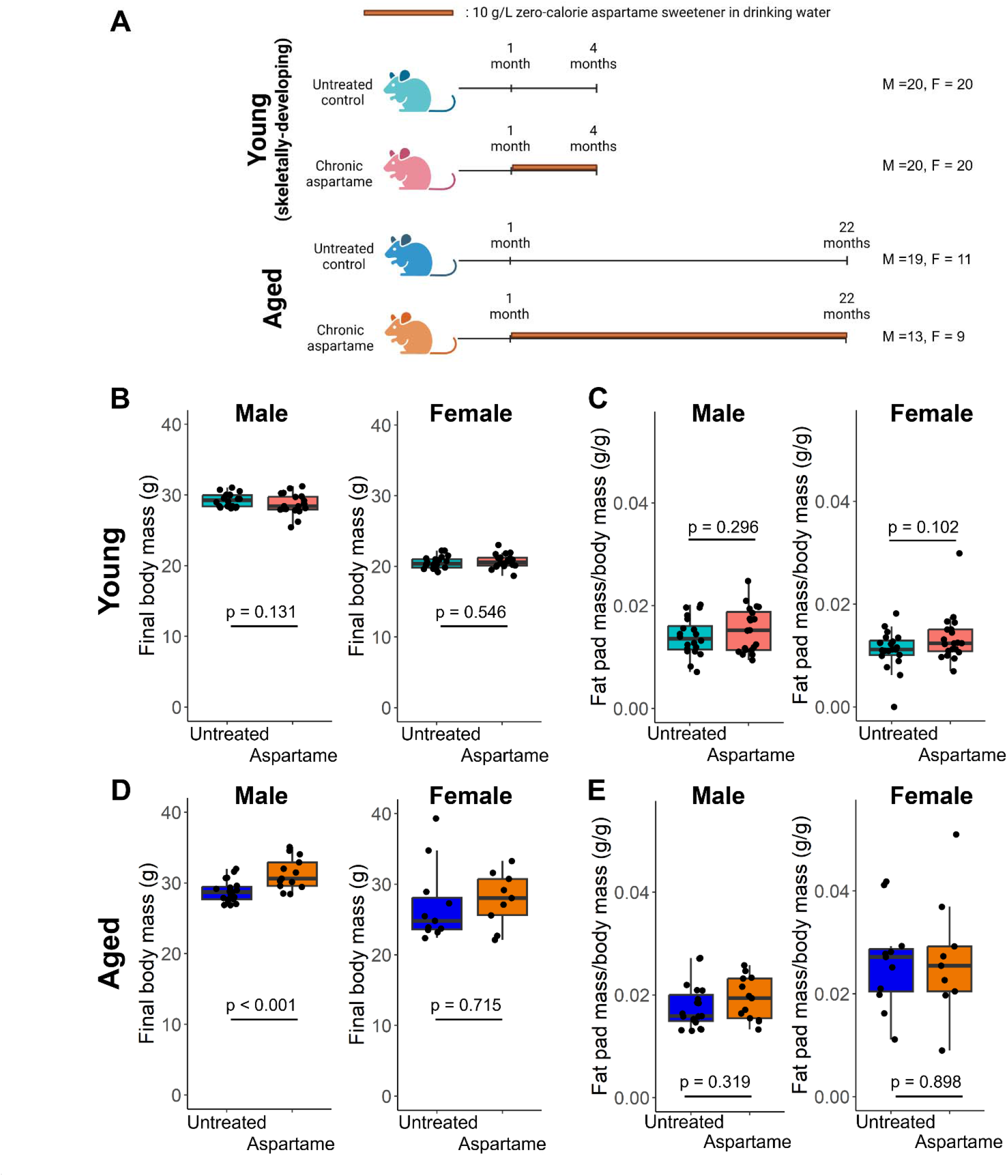
Aged male mice receiving aspartame had greater body mass. A) Study design. B-C) Aspartame treatment to mice in the young cohort had no effect on body mass or fat pad mass. D) Body mass at euthanasia was greater in aged males chronically treated with aspartame but not aged females. E) Aspartame treatment did not influence perigonadal fat pad mass in aged mice. A one-way ANOVA was used to calculate differences in body mass and normalized fat pad mass between treatment groups in each age and sex.

### Geometric measurements of femoral cortical bone

The strength of bone is influenced by both geometry and the material properties of the bone matrix. The size and geometric properties of each femur must first be measured using micro-computed X-ray tomography as follows: femora were thawed to room temperature and scanned using 10-micron voxel size resolution with a mineral calibration phantom (Bruker SkyScan 1276 mouse CT, Bruker Corporation, Billerica, MA, US). Images were processed using a standard methodology and geometric measurements were made from the diaphyseal cross-section in the microcomputed tomography images (9,15). Images were analyzed using a custom Java code and run in Fiji (v.2.3.051) with a BoneJ plug-in. Briefly, a Gaussian blur was applied (σ = 1), the mid-slice of the mid-diaphysis was determined, and a threshold was applied using the Otsu method (16,17). Cortical cross-sectional area, cortical thickness, moment of inertia (I) about the medial-lateral axis (direction of loading), distance from the neutral axis to the edge of the bone surface (c), and section modulus (I/c) were measured.

### Biomechanics

The mechanical properties of the mouse femur were determined by applying three point bending until failure. Three point bending tests in this manner directly provide the whole bone strength (maximum load the bone can experience in bending). Whole bone strength and measures of bone cross-sectional geometry can be used to calculate bone tissue strength (a mechanical property of the bone matrix itself, that is independent of bone size or geometry). Mechanical testing in three point bending was performed as follows: the right femur was thawed, hydrated with phosphate buffered saline and tested using three-point bending in anterior-to-posterior direction to failure (loading rate = 0.1 mm/sec, span length = 7.5 mm between outer loading pins) using a materials testing device (858 Mini Bionix; MTS, Eden Prairie, MN, US). Force versus displacement curves were collected using a 11.34 kg (25 lbs) load cell (MLP-25, Transducer Techniques, Temecula, CA, US; weekly manual calibrations) and linear variable differential transducer at 100-Hz sampling. Data was processed using a custom script in Matlab (R2021b, MathWorks, Natick, MA, US) using a smoothing function to minimize noise. The maximum breaking force and displacement were recorded. A 10% reduction in slope method was used to identify the yield force and yield displacement (18). The maximum breaking force was used to calculate whole bone strength (max force*span length/4). Bone tissue strength was calculated using whole bone strength and geometric measurements from micro-computed X-ray tomography (whole bone strength/section modulus). Whole bone strength is determined by the size/geometry of the bone cross-section at the point of loading as well as the strength of the bone matrix itself whereas bone tissue strength is determined by the strength of the bone matrix (independent of bone size or geometry).

### *16S* rRNA gene sequencing analysis

16S rRNA gene sequencing was used to determine the composition of the microbiota from fecal samples collected at 4 months (young cohort) or 22 months of age (aged cohort). The University of California San Diego Microbiome Core carried out DNA extraction, purification, library preparation, and sequencing. DNA was isolated and purified using a liquid handler robot (MagMAX Microbiome Ultra Nucleic Acid Isolation Kit, Thermo Fisher Scientific, US). For quality control, blanks and mock communities were used throughout the processing pipeline (Zymo Research Corporation, US). 16S rRNA gene amplification was completed using a protocol from the Earth Microbiome Project (19). Unique forward primer barcodes (Illumina) were used to amplify the V4 region of the 16S rRNA gene (515fB-806r; forward – GTGYCAGCMGCCGCGGTAA, reverse – GGACTACNVGGGTWTCTAAT; (20,21)).

Amplification was individually carried out on each sample as a single reaction (94°C 3 min, 94°C 45 sec x35, 50°C 60 sec x35, 72°C 90 sec x35, 72°C 10 min, 4°C hold), equal volumes of each amplicon were pooled, and libraries were sequenced on Illumina MiSeq using paired-end 150 bp cycles (22). Quality control trimming and taxonomic classification was carried out using QIIME2 (v. 2020.6) with a SILVA database (SSU r138-1) (23). Reads were assigned at the genus level and normalized with a single rarefaction step (feature count cut-off 46067; range of feature sizes in samples: 46068-233105; average and standard deviation of feature size in samples: 110390 ± 59424) (24). Feature count cut-off was determined using standard rarefaction methods in an effort to maximize the number of features relative to the percent of samples retained (12,24,25). Alpha (Shannon index) and beta diversity (Bray-Curtis dissimilarity) were calculated using the amplicon sequence variant (ASV) table from QIIME2 output using the vegan package (v. 2.5-7) in RStudio (12,26). The Bray-Curtis dissimilarity matrix was rarefied and a principal coordinate analysis (PCoA) was carried out by sex and treatment group. Microbiome Multivariate Associations with Linear Models (MaAsLin2) was carried out to identify univariate associations between treatment groups, sex, and microbial abundance (27). Specifically, genera-level taxa were used in the MaAsLin2 analysis that had > 10% prevalence and post-hoc q-values associating treatment groups and microbes were calculated with the Benjamini-Hochberg method (27). A secondary differential abundance analysis (ALDEx2 R package) was performed using a rarefied ASV table (separated by sex) to create Monte Carlo simulations of Dirichlet distributions of each sample (28,29). The distributions were transformed using a centered log ratio transform and a generalized linear model was generated based on treatment (aspartame versus untreated) (29). Volcano plots were generated by sex and significant ASVs (p < 0.05; magnitude of fold change > |1|) are shown in upper left and right corners.

### Statistical analysis

Values are shown as the mean ± standard deviation. Statistical analyses were performed using RStudio (v. 1.4.1106, Boston, MA, US, 2021) (30). One-way analysis of variance (ANOVA) was used to calculate significance between treatment groups by age and sex in body mass, fat pad mass, bone biomechanics, femoral geometry, alpha diversity, and microbial abundance. For parameters that were correlated (section modulus and whole bone strength), an analysis of covariance (ANCOVA) was performed to detect differences in whole bone strength between groups when accounting for differences in section modulus (9). Permutational multivariate analysis of variance (PERMANOVA; adonis2 function) was used to calculate differences in beta diversity (microbiota composition) by treatment group (31). A Pearson’s correlation matrix between femoral geometry, body mass, fat pad mass, and biomechanics was performed (95% confidence intervals are shown; p-values < 0.05 considered significant). Femoral geometry and bone biomechanics are shown without (main figure) and with normalization (supplementary table) by body mass. Normalization by body mass was performed using a regression-based approach (32).

## Results

### Aged male mice receiving aspartame had greater body mass

Aged male mice (22 month) receiving chronic high dose aspartame (10 g/L) had greater body mass (**Figure 1D**; p < 0.001) and comparable perigonadal fat pad mass (normalized by body mass; **Figure 1E**; p = 0.319) relative to untreated mice. Young (4 month) males and females (both young and aged) receiving aspartame did not have differences in body mass or fat pad mass relative to untreated mice.

### Aged male mice receiving aspartame had greater whole bone strength and femoral geometry

Aged male mice treated with aspartame had greater whole bone strength (**Figure 2E**; p = 0.006), maximum load (**Table 1**; p = 0.006), and work to failure (**Table 1**; p = 0.028) relative to mice not receiving aspartame. Additionally, aspartame treated aged male mice had greater femoral geometry including section modulus (**Figure 2F**; p < 0.001), cross-sectional area (**Figure 2G**; p = 0.021), and moment of inertia (**Figure 2H**; p < 0.001) than untreated mice. Differences in whole bone strength and femoral geometry were not detected in young males (4 month) or in females at either age. There were no differences in femur length, yield load, or yield displacement at either timepoint in either sex (**Table 1**).

**Figure 2:**
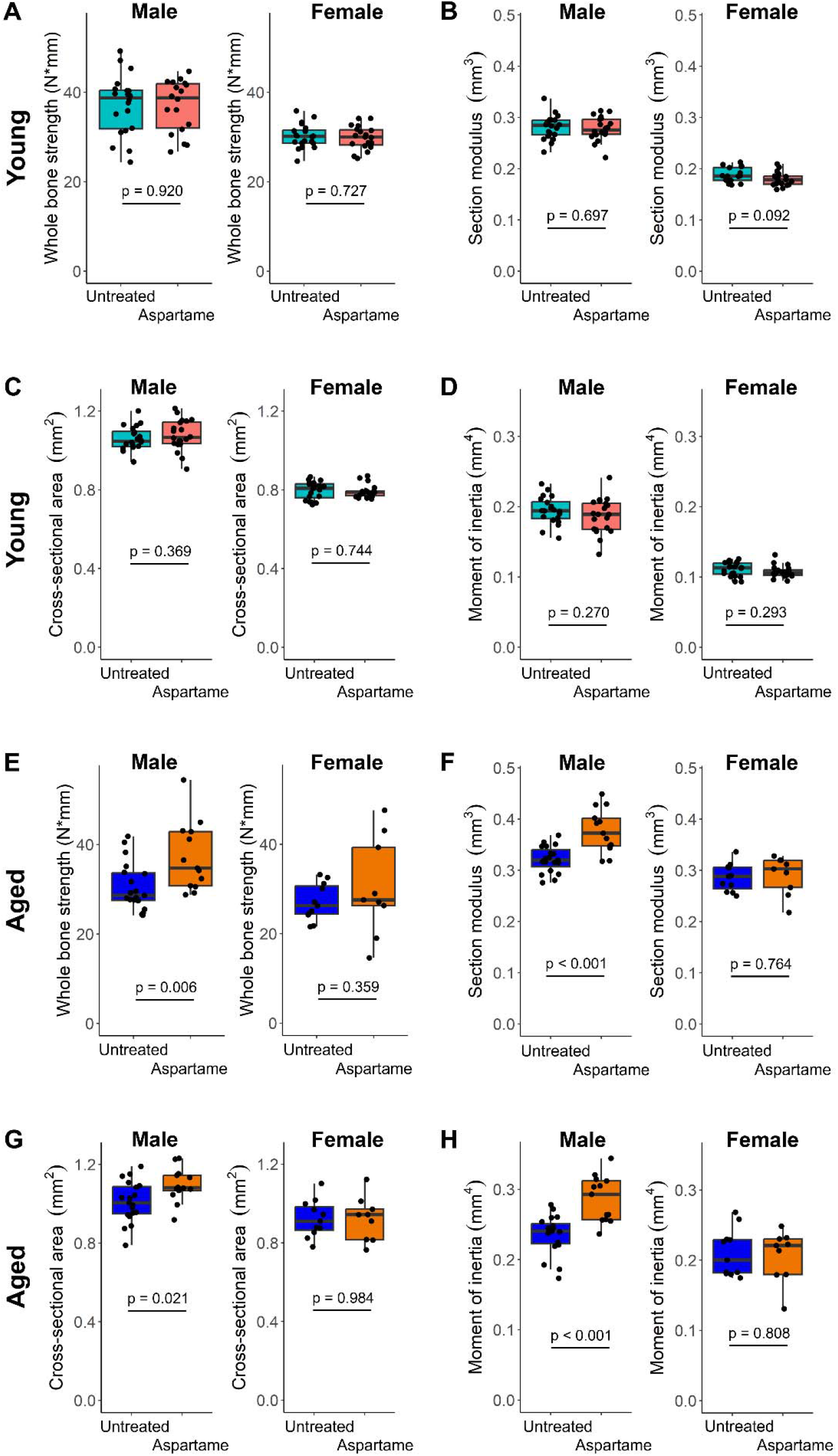
Aged male mice treated with aspartame had greater whole bone strength and femoral geometry. A-D) Aspartame treatment to mice in the young cohort had no effect on whole bone strength, section modulus, cross-sectional area, or moment of inertia. E-H) Whole bone strength, section modulus, cross-sectional area, and moment of inertia were greater in aged males treated with aspartame but not females. A one-way ANOVA was used to calculate differences in biomechanics and femoral geometry between treatment groups in each age and sex.

**Table 1:** Measurements of bone geometry and biomechanics are shown. Statistical comparisons are only made within sex and age groups. Results are shown as the mean ± standard deviation. Measurements normalized for differences in body mass are shown in **Supplementary** Table 3.

| Treatment Group | Femur length (mm) | Tissue strength (N/mm<sup>2</sup>) | Distance to neutral axis (mm) | Maximum load (N) | Yield displacement (mm) | Stiffness (N/mm) | Yield load (N) | Work to failure (N*mm) |
| --- | --- | --- | --- | --- | --- | --- | --- | --- |
| *Males – Aged (22 months)* |  |  |  |  |  |  |  |  |
| Chronic aspartame | 16.394 ± 0.285<br>p=0.794 | 99.216 ± 20.251<br>p=0.551 | 0.759 ± 0.024<br>p=0.071 | 19.847 ± 4.072<br>p=0.006 | 0.127 ± 0.034<br>p=0.679 | 100.290 ± 18.198<br>p=0.516 | 11.334 ± 2.556<br>p=0.594 | 7.925 ± 2.714<br>p=0.028 |
| Unaltered | 16.361 ± 0.381 | 95.315 ± 16.254 | 0.733 ± 0.046 | 16.275 ± 2.834 | 0.134 ± 0.047 | 95.237 ± 23.208 | 10.827 ± 2.602 | 5.485 ± 3.080 |
| *Males – Young (4 months)* |  |  |  |  |  |  |  |  |
| Chronic aspartame | 16.003 ± 0.332<br>p=0.127 | 132.370 ± 25.873<br>p=0.979 | 0.669 ± 0.052<br>p=0.093 | 19.509 ± 3.116<br>p=0.961 | 0.146 ± 0.028<br>p=0.382 | 131.123 ± 19.389<br>p=0.386 | 15.613 ± 2.646<br>p=0.795 | 5.162 ± 3.559<br>p=0.868 |
| Unaltered | 16.119 ± 0.190 | 132.586 ± 23.326 | 0.692 ± 0.026 | 19.561 ± 3.475 | 0.135 ± 0.047 | 136.482 ± 18.733 | 15.313 ± 4.030 | 4.955 ± 4.146 |
| *Females – Aged (22 months)* |  |  |  |  |  |  |  |  |
| Chronic aspartame | 16.111 ± 0.256<br>p=0.604 | 103.754 ± 31.899<br>p=0.422 | 0.706 ± 0.051<br>p=0.325 | 16.212 ± 5.821<br>p=0.359 | 0.138 ± 0.055<br>p=0.935 | 105.396 ± 35.614<br>p=0.845 | 12.019 ± 6.046<br>p=0.770 | 5.935 ± 3.203<br>p=0.034 |
| Unaltered | 16.187 ± 0.365 | 94.917 ± 14.656 | 0.732 ± 0.060 | 14.429 ± 2.213 | 0.137 ± 0.025 | 102.846 ± 21.369 | 12.618 ± 2.007 | 2.997 ± 2.306 |
| *Females – Young (4 months)* |  |  |  |  |  |  |  |  |
| Chronic aspartame | 15.467 ± 0.272<br>p=0.560 | 166.541 ± 14.613<br>p=0.185 | 0.598 ± 0.029<br>p=0.441 | 15.833 ± 1.354<br>p=0.647 | 0.131 ± 0.040<br>p=0.094 | 97.389 ± 12.831<br>p=0.001 | 10.595 ± 1.777<br>p=0.748 | 4.241 ± 2.579<br>p=0.968 |
| Unaltered | 15.509 ± 0.248 | 159.252 ± 17.276 | 0.591 ± 0.030 | 15.798 ± 1.770 | 0.111 ± 0.032 | 111.562 ± 12.687 | 10.406 ± 1.876 | 4.275 ± 2.726 |

When normalized by body mass, differences in cross-sectional area and whole bone strength between aged males with and without aspartame were not significantly different (**Supplementary Table 3**) (32). Femoral geometry (cross-sectional area, moment of inertia, and section modulus) was positively correlated with body mass and fat pad mass in both aged males (**Supplementary Table 4**) and females (**Supplementary Table 5**). Additionally, cross-sectional area, moment of inertia, and section modulus were positively correlated with whole bone strength in both sexes in aged mice.

The biomechanical analysis from the young (4 month) male and female mice suggested that chronic aspartame did not affect bone strength or geometry in either sex during early growth. No differences in biomechanical parameters were associated with aspartame treatment after normalizing by body mass.

### Aged male and female mice receiving aspartame had a different microbiota than untreated mice

Since we did not observe the difference in whole bone strength in the young mouse cohort as we had observed in our prior work, we looked in greater detail at changes in the gut microbiota associated with aspartame in the current study. Both aged males and females receiving chronic aspartame had a significantly different microbiota composition (Bray-Curtis beta diversity) relative to untreated (M: p = 0.007; F: p = 0.026) (**Figure 3A**, **Supplementary** Figure 1). The differences in beta diversity in aged males receiving aspartame appeared to be caused by a few individual mice that had a drastically different taxonomic abundance of microbes compared to untreated mice (**Figure 3B**). Aspartame treatment did not affect alpha diversity (**Supplementary** Figure 2: observed richness, Shannon diversity) in either aged males or females. MaAsLin analysis found that *Odoribacter* was significantly decreased in aged males and females receiving aspartame relative to untreated mice (**Figure 3C**; M: p = 0.002; F: p = 0.052). Differential abundance analysis by ALDEx2 determined that 1.8% (15/852) and 2.4% (21/875) of the ASVs identified in aged males and females were significant (p < 0.05; magnitude of fold change > |1|) in mice that received aspartame relative to untreated mice (**Figure 3D**). Notably, all the significant ASVs identified in aged males and females belonged to either *Bacteroidetes* or *Firmicutes* phyla (**Supplementary Tables 6**-**7**).

**Figure 3:**
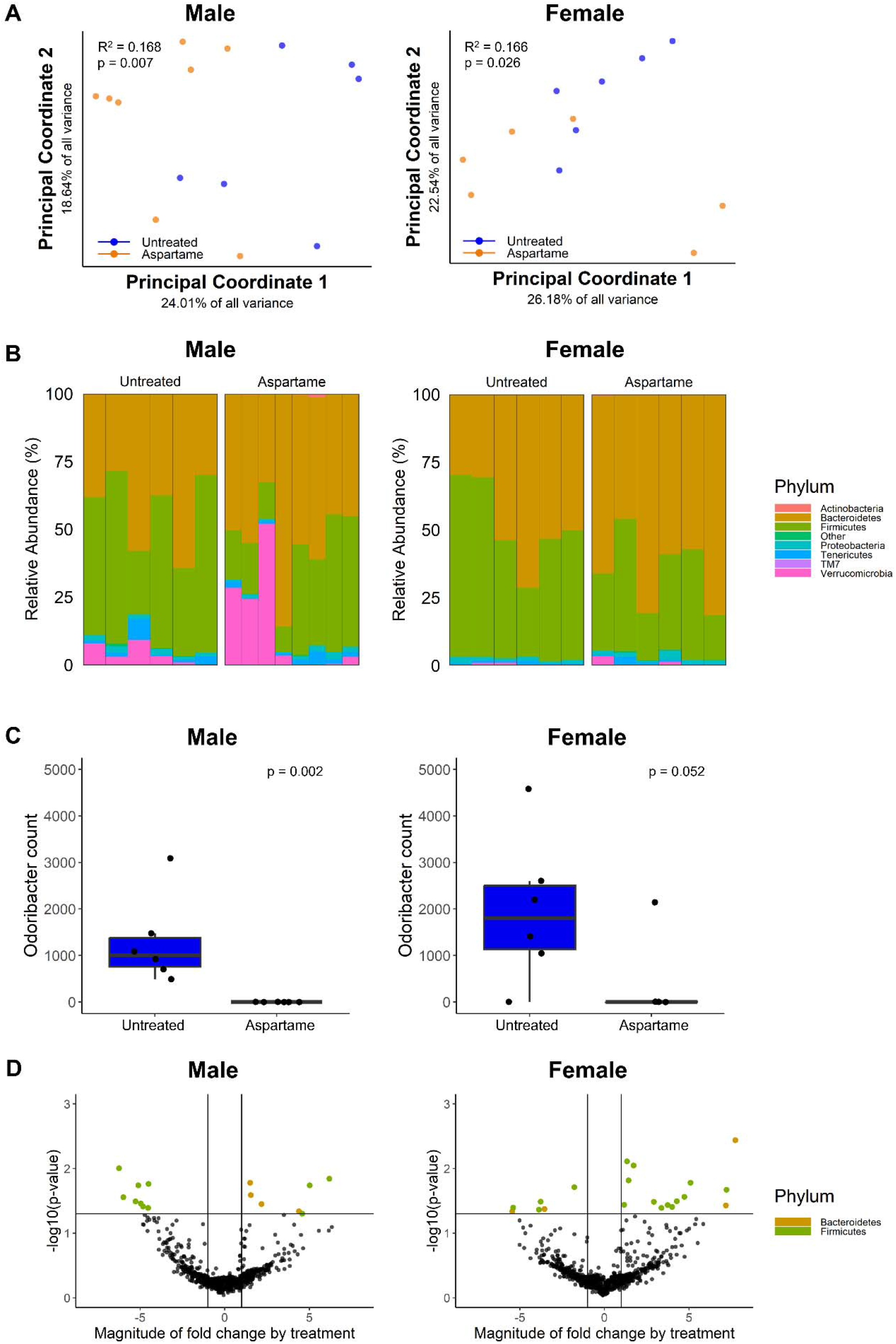
Aged male and female mice receiving chronic aspartame had a different microbiota than untreated mice. A) Both aged males and females receiving aspartame had a different composition of the microbiota (Bray-Curtis beta diversity) than untreated mice. B) A subset of aged male mice receiving aspartame had a drastically different relative abundance of taxa compared to untreated mice. C) Aspartame treatment decreased the abundance of *Odoribacter* in both aged males and females relative to untreated. D) Differential abundance analysis by ALDEx2 identified significant ASVs from the *Bacteroidetes* and *Firmicutes* phyla in both males and females.

## Discussion

In this study, we evaluated the effects of long-term high dose aspartame on body mass, whole bone strength, femoral geometry, and microbiota composition in a young (4 month old) and aged (22 month old) cohort of male and female mice. Whole bone strength and section modulus were both increased in aged males that were treated with aspartame. However, there were no differences in bone tissue strength (calculated from the whole bone strength/section modulus) between aged groups with and without aspartame. Further, whole bone strength and femoral geometry in aged males receiving aspartame were not significantly different relative to untreated mice when normalized by body mass and femoral geometry was positively correlated with body mass and whole bone strength. Collectively, these findings suggested that the increase in whole bone strength in the aged males was primarily determined by the greater femoral geometry due to a greater body mass in mice receiving aspartame, rather than tissue-level changes to bone (33,34). Bone size is strongly correlated with body mass in mice (35).

Thirty one percent of the aged females (22 month) treated with aspartame in the study died prior to the end of the study (deaths occurred between 9-22 months of age) and it is plausible that the decrease in sample size and power may partially explain why the effect on bone mechanical properties and femoral geometry was not detected in aged females. Staff veterinarians did not determine the cause of the premature death of the aged females treated with aspartame. In contrast, aged males treated with aspartame had 100% survivorship.

The composition of the microbiota differed in both aged males and females receiving aspartame relative to untreated mice with a decreased abundance of *Odoribacter* in mice that received aspartame. *Odoribacter* is a member of *Bacteroidetes* phylum and *Porphyromondaceae* family that has previously been shown to be decreased with aspartame treatment (7). Additionally, human-derived isolates of *Odoribacter sp.* have been used as probiotics to consume excess succinate to modulate glucose tolerance and inflammation in mice (36) and decreased abundance of *Odoribacter* with aspartame may partially explain a greater body mass in aged male mice (37). The secondary differential abundance analysis via ALDEx2 was confirmatory of the results from MaAsLin2 and identified significant ASVs in aged males and females from the *Bacteroides* phylum and *Porphyromondaceae* family.

In young mice (4 month age), aspartame treatment did not influence body mass, demonstrating that long-term exposure to aspartame (> 3 months) had greater effects on body mass than exposure in the short-term (< 3 months). Our findings were consistent with a prior study in which chronic aspartame impaired the animal’s neurometabolic response to appropriately sense sweet-taste resulting in increased food consumption and weight gain; an effect that was more pronounced in males (38). However, our findings were inconsistent with our prior aspartame study that was performed in a small cohort of young skeletally developing male mice (4 months of age). Our prior aspartame study resulted in an increase in whole bone strength (independent of the geometry of the bone) that was not observed in the current, more comprehensive follow-up study. It is unclear why we observed these differences in aspartame influencing bone strength in young mice across independent studies, however it could potentially be related to skeletal differences in the breeders, cage activity, environmental factors, changes in the manufacturing process of aspartame sweetener, different starting microbiota composition, or other confounding variables that have yet to be determined. For this reason, it is important to replicate studies of the microbiota using the same stimulus to confirm the consistency of the results. When comparing the Bray-Curtis beta diversity of the microbiota across our two independent studies it appeared that in our initial study (Study 1) the aspartame resulted in a different microbiota than in the untreated mice (p = 0.002), but in our follow-up study (current; Study 2) the aspartame treated mice had a microbiota more similar to the unaltered mice (p = 0.031) (**Supplementary** Figure 3). It is possible that mice had different microbiota at the start of each of the two studies and therefore the aspartame had a different effect on the microbiota, however, further analysis would be needed to confirm this.

This study has several limitations related to its scope. Specifically, our study was not designed to explain the mechanism of how the increased body mass caused by aspartame increased femoral geometry and whole bone strength in aged male mice. It is conceivable that aspartame increased mouse cage activity leading to increased loading and bone formation. It has also been reported that chronic aspartame impairs the function of intestinal alkaline phosphatase through its byproduct phenylalanine causing translocation of gut microbes and inflammation (39,40).

Therefore, it is also possible that aspartame could have inactivated intestinal alkaline phosphatase and interfered with bone metabolism signaling downstream (TLR4/NF-ĸB signaling pathways) (39) and/or microbial metabolites or translocated microbes were directly influencing bone metabolism. Future work involves analysis of microbial-derived metabolites in mice receiving aspartame. Further, our study was limited in that it did not measure food intake, which could provide more insight into why aspartame treated aged male mice had an increased body mass.

Our study involved mice receiving an extraordinarily high dose of aspartame (estimated daily intake 1500 mg/kg, assuming 2-3 mL water consumed/day, human equivalent dose 122 mg/kg) (41) more than a reasonable dose that would be consumed by humans(acceptable daily intake = 50 mg/kg body mass) (42). However, our dose was within the range in prior carcinogenicity studies in rodents and accounted for differences in the rate of aspartame metabolism in mice versus humans (42,43). Our goal was to evaluate if aspartame influenced the microbiota and if those differences in the microbiota were associated with bone strength in young and aged mice under high dose aspartame conditions.

In summary, aged male mice receiving chronic aspartame had greater body mass, whole bone strength, and femoral geometry relative to untreated mice; an effect that was not observed in aged females. The greater whole bone strength in aged male mice receiving aspartame appeared to have been related to greater animal body mass and not to changes in bone tissue strength.

Aspartame treatment had a similar effect on the composition of the gut microbiota and decreased *Odoribacter* in both aged males and females, demonstrating that aspartame did not have a sex-dependent effect on the microbiota. Across independent studies young male mice receiving aspartame did not have consistent effects on whole bone strength or microbiota composition. Our study highlights the need for replication in microbiome studies and the importance of long-term rodent studies to understand the effects of compounds of interest across the lifespan.

## Acknowledgements

Dr. Teresa Porri for acquisition of micro-CT imaging data of right femora. Imaging data was acquired through the Cornell Institute of Biotechnology’s Imaging Facility, with NIH S10OD2549 for the SkyScan 1276 mouse CT.

## Abbreviations

ALDEx: = ANOVA-like differential expression tool
ANCOVA: = analysis of covariance
ANOVA: = analysis of variance
ASV: = amplicon sequence variant
MaAsLin: = microbiome multivariate associations with linear models
NF-ĸB: = nuclear factor kappa light chain enhancer of activated B cells
PCoA: = principal coordinate analysis
PERMANOVA: = permutational multivariate analysis of variance
TLR4: = toll-like receptor 4

## Statement of authors’ contributions to manuscript

E.L.C, I.L.B, S.L.B, and C.J.H. designed research; E.L.C, C.L., A.L.M., J.C.N., E.B., M.G. conducted research; E.L.C., C.L., A.L.M., N.C., P.J.T., I.L.B., S.L.B., and C.J.H. analyzed data; and E.L.C., P.J.T., I.L.B., S.L.B., and C.J.H. wrote the paper. C.J.H. had primary responsibility for final content. All authors read and approved the final manuscript.

## Declaration of generative AI in scientific writing

The authors did not use any AI or AI-assisted technologies in the writing of this manuscript.

---

**Supplementary Figure 1:**
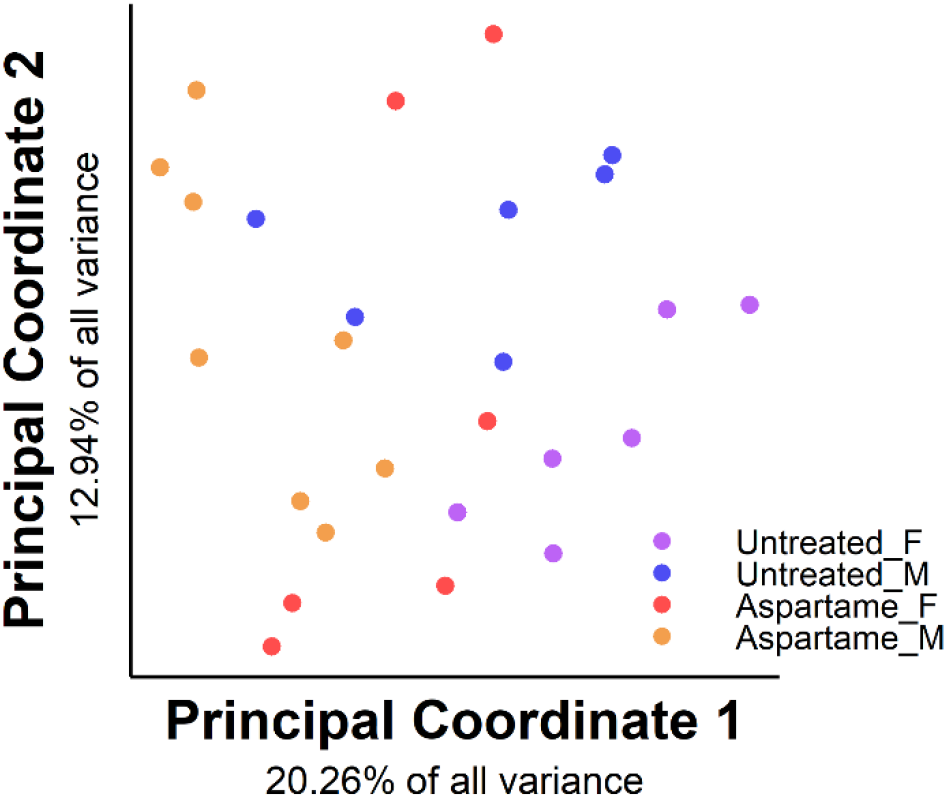
Combined aged male and female aspartame treated and untreated mice Bray-Curtis beta diversity.

**Supplementary Figure 2:**
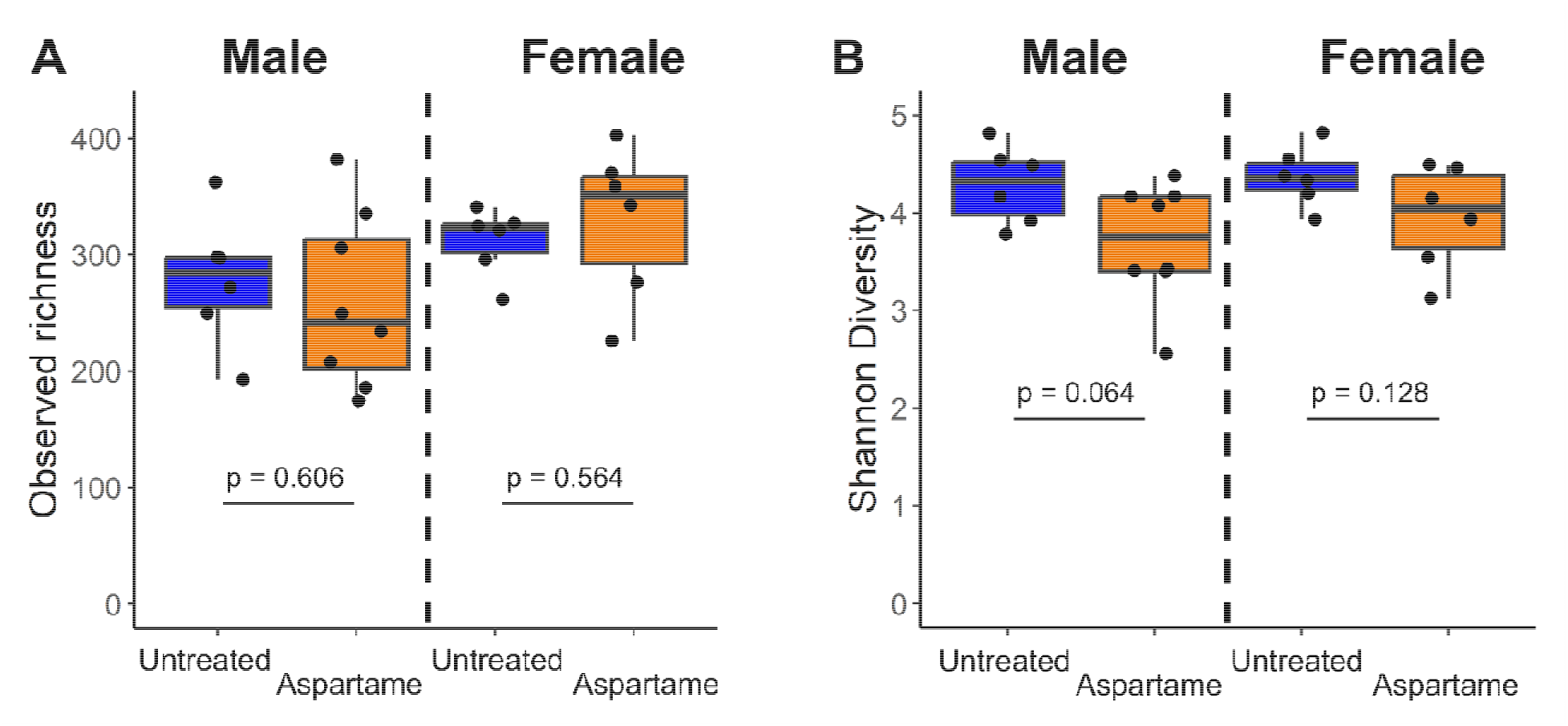
Aspartame treatment did not influence alpha diversity in aged males or females. A-B) Aspartame treatment did not affect observed richness or Shannon diversity in aged males or females.

**Supplementary Figure 3:**
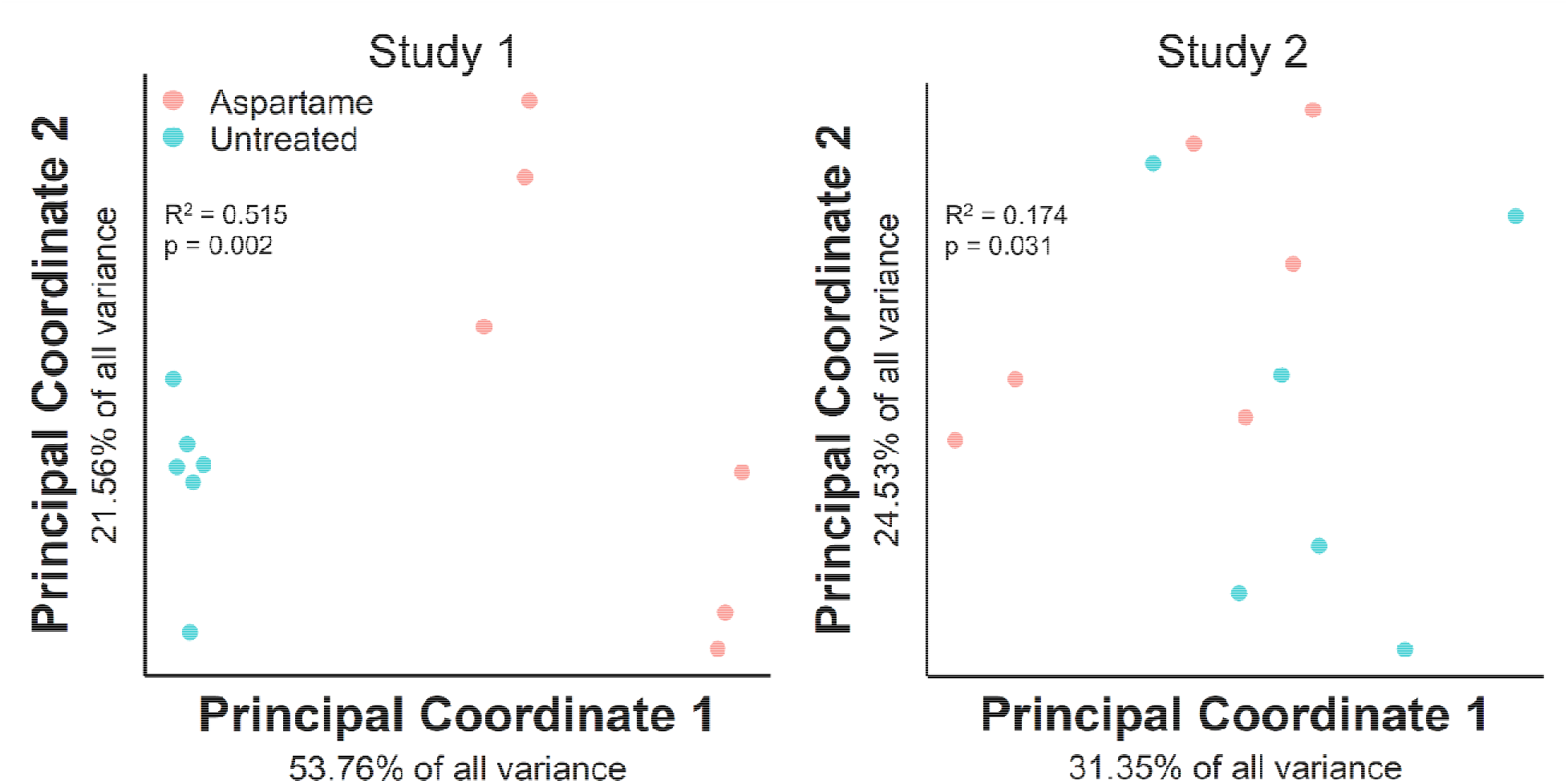
Across two independent studies in male mice treated with aspartame from 1-4 months of age, aspartame had a different effect on the microbiota (Bray-Curtis beta diversity). In our initial study (Study 1 (9)), male mice receiving aspartame had a large difference in the composition of the microbiota relative to untreated mice (p = 0.002). In our current follow-up study (Study 2), male mice receiving aspartame had a microbiota composition more similar to that of the unaltered mice relative to Study 1 (p = 0.031).

**Supplementary Table 1:**
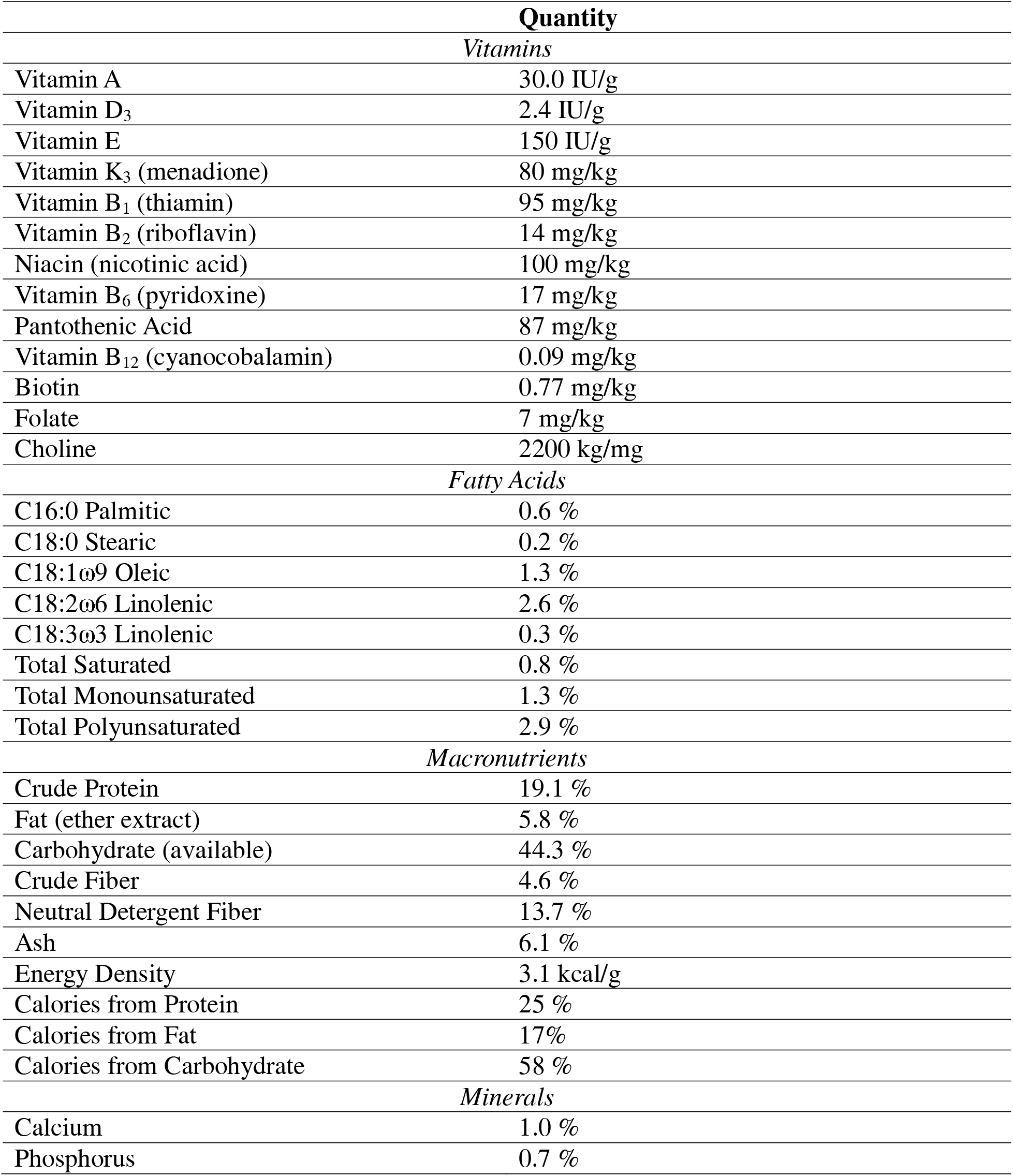
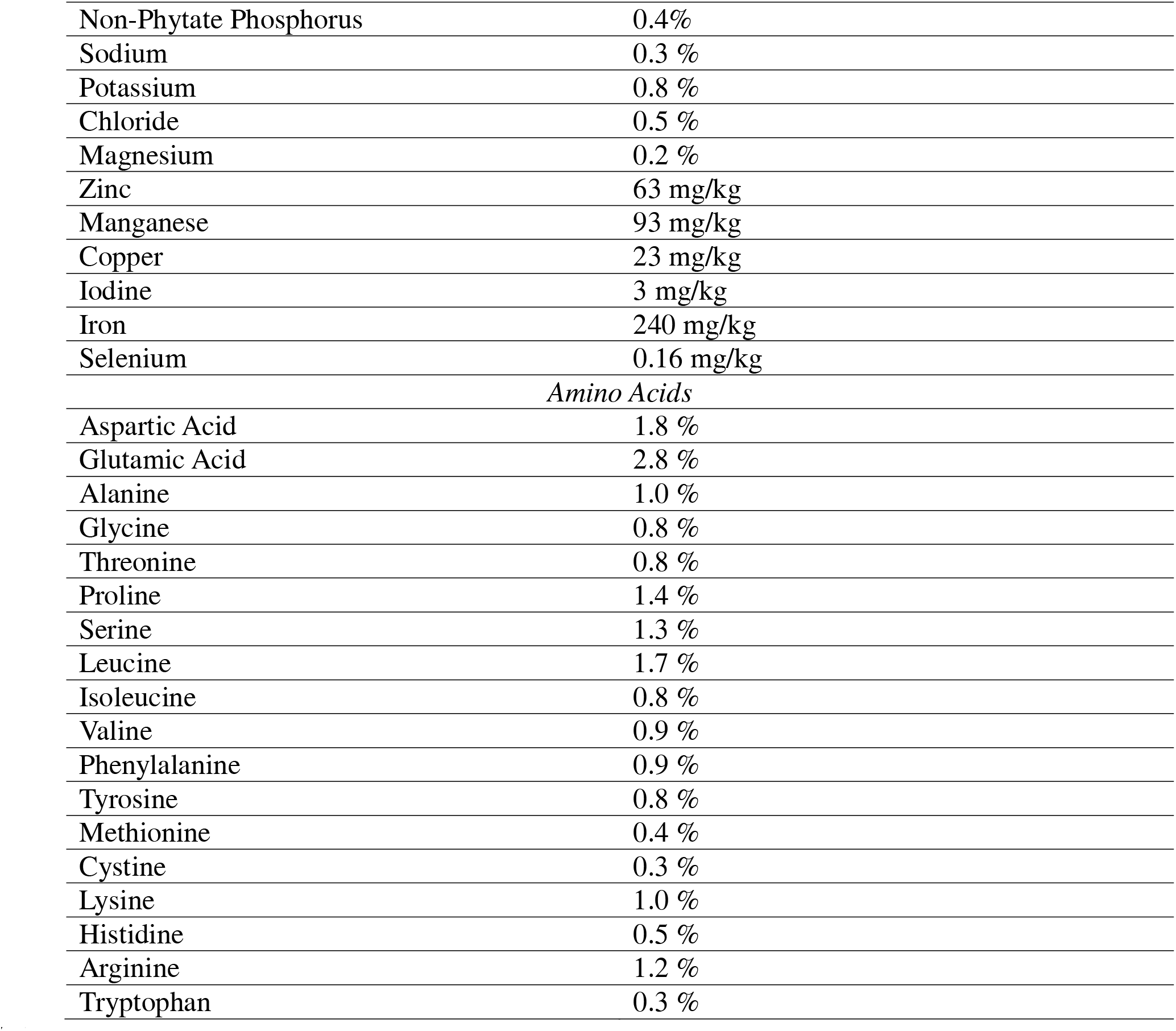
Ingredient list of Teklad LM-485 sterilizable mouse/rat chow consumed by all mice in the study (with and without aspartame treatment in drinking water) as provided by the manufacturer.

|  | Quantity |
| --- | --- |
| *Vitamins* |  |
| Vitamin A | 30.0 IU/g |
| Vitamin D<sub>3</sub> | 2.4 IU/g |
| Vitamin E | 150 IU/g |
| Vitamin K<sub>3</sub> (menadione) | 80 mg/kg |
| Vitamin B<sub>1</sub> (thiamin) | 95 mg/kg |
| Vitamin B<sub>2</sub> (riboflavin) | 14 mg/kg |
| Niacin (nicotinic acid) | 100 mg/kg |
| Vitamin B<sub>6</sub> (pyridoxine) | 17 mg/kg |
| Pantothenic Acid | 87 mg/kg |
| Vitamin B<sub>12</sub> (cyanocobalamin) | 0.09 mg/kg |
| Biotin | 0.77 mg/kg |
| Folate | 7 mg/kg |
| Choline | 2200 mg/kg |
| *Fatty Acids* |  |
| C16:0 Palmitic | 0.6 % |
| C18:0 Stearic | 0.2 % |
| C18:1ω9 Oleic | 1.3 % |
| C18:2ω6 Linolenic | 2.6 % |
| C18:3ω3 Linolenic | 0.3 % |
| Total Saturated | 0.8 % |
| Total Monounsaturated | 1.3 % |
| Total Polyunsaturated | 2.9 % |
| *Macronutrients* |  |
| Crude Protein | 19.1 % |
| Fat (ether extract) | 5.8 % |
| Carbohydrate (available) | 44.3 % |
| Crude Fiber | 4.6 % |
| Neutral Detergent Fiber | 13.7 % |
| Ash | 6.1 % |
| Energy Density | 3.1 kcal/g |
| Calories from Protein | 25 % |
| Calories from Fat | 17% |
| Calories from Carbohydrate | 58 % |
| *Minerals* |  |
| Calcium | 1.0 % |
| Phosphorus | 0.7 % |
| Non-Phytate Phosphorus | 0.4% |
| Sodium | 0.3 % |
| Potassium | 0.8 % |
| Chloride | 0.5 % |
| Magnesium | 0.2 % |
| Zinc | 63 mg/kg |
| Manganese | 93 mg/kg |
| Copper | 23 mg/kg |
| Iodine | 3 mg/kg |
| Iron | 240 mg/kg |
| Selenium | 0.16 mg/kg |
| *Amino Acids* |  |
| Aspartic Acid | 1.8 % |
| Glutamic Acid | 2.8 % |
| Alanine | 1.0 % |
| Glycine | 0.8 % |
| Threonine | 0.8 % |
| Proline | 1.4 % |
| Serine | 1.3 % |
| Leucine | 1.7 % |
| Isoleucine | 0.8 % |
| Valine | 0.9 % |
| Phenylalanine | 0.9 % |
| Tyrosine | 0.8 % |
| Methionine | 0.4 % |
| Cystine | 0.3 % |
| Lysine | 1.0 % |
| Histidine | 0.5 % |
| Arginine | 1.2 % |
| Tryptophan | 0.3 % |

**Supplementary Table 2:** Sample sizes included in each treatment group.

| Treatment Group | Sample Size |
| --- | --- |
| *Aged (22 months)* |  |
| Chronic aspartame | 13 (M); 9 (F) |
| Untreated | 19 (M); 11 (F) |
| *Young (4 months)* |  |
| Chronic aspartame | 20 (M); 20 (F) |
| Untreated | 20 (M); 20 (F) |

**Supplementary Table 3:** Biomechanics and femoral geometry parameters normalized by body mass. The mean ± standard deviation are provided for each parameter.

| Treatment Group | Femur length (mm) | Whole bone strength (N*mm) | Cross-sectional area (mm<sup>2</sup>) | Moment of inertia (mm<sup>4</sup>) | Section modulus (mm<sup>3</sup>) | Tissue strength (N/mm<sup>2</sup>) | Distance to neutral axis (mm) |
| --- | --- | --- | --- | --- | --- | --- | --- |
| *Males – Aged (22 months)* |  |  |  |  |  |  |  |
| Chronic aspartame | 16.377 ± 0.284<br>p=0.859 | 36.435 ± 7.543<br>p=0.051 | 1.066 ± 0.073<br>p=0.655 | 0.273 ± 0.025<br>p=0.004 | 0.359 ± 0.032<br>p=0.004 | 101.883 ± 21.223<br>p=0.469 | 0.760 ± 0.024<br>p=0.187 |
| Untreated | 16.354 ± 0.383 | 31.994 ± 4.823 | 1.053 ± 0.080 | 0.246 ± 0.023 | 0.331 ± 0.020 | 97.043 ± 16.094 | 0.741 ± 0.044 |
| *Males – Young (4 months)* |  |  |  |  |  |  |  |
| Chronic aspartame | 16.033 ± 0.230<br>p=0.156 | 37.013 ± 5.824<br>p=0.879 | 1.078 ± 0.076<br>p=0.174 | 0.190 ± 0.020<br>p=0.635 | 0.281 ± 0.020<br>p=0.739 | 131.618 ± 25.027<br>p=0.955 | 0.675 ± 0.043<br>p=0.174 |
| Untreated | 16.130 ± 0.188 | 36.700 ± 6.624 | 1.049 ± 0.055 | 0.193 ± 0.018 | 0.279 ± 0.020 | 132.065 ± 23.878 | 0.691 ± 0.026 |
| *Females – Aged (22 months)* |  |  |  |  |  |  |  |
| Chronic aspartame | 16.107 ± 0.253<br>p=0.552 | 29.845 ± 9.778<br>p=0.727 | 0.914 ± 0.085<br>p=0.684 | 0.203 ± 0.022<br>p=0.414 | 0.288 ± 0.020<br>p=0.961 | 102.880 ± 30.967<br>p=0.736 | 0.703 ± 0.041<br>p=0.121 |
| Untreated | 16.192 ± 0.357 | 28.669 ± 4.620 | 0.928 ± 0.064 | 0.212 ± 0.024 | 0.288 ± 0.022 | 99.373 ± 12.941 | 0.735 ± 0.045 |
| *Females – Young (4 months)* |  |  |  |  |  |  |  |
| Chronic aspartame | 15.470 ± 0.224<br>p=0.372 | 29.958 ± 2.616<br>p=0.664 | 0.794 ± 0.030<br>p=0.685 | 0.108 ± 0.008<br>p=0.272 | 0.180 ± 0.012<br>p=0.073 | 166.684 ± 15.022<br>p=0.206 | 0.598 ± 0.028<br>p=0.406 |
| Untreated | 15.529 ± 0.182 | 30.307 ± 2.363 | 0.798 ± 0.041 | 0.111 ± 0.010 | 0.189 ± 0.014 | 159.751 ± 16.744 | 0.590 ± 0.029 |

**Supplementary Table 4:** Pearson correlations between body mass, fat pad mass, biomechanics, and femoral geometry in aged male mice. The upper right shows r values while the bottom left shows confidence intervals (95%) and p-values are shown (bold and underlined: p < 0.05).

|  | Body mass (g) | Fat pad mass (g) | Cross-sectional area (mm<sup>2</sup>) | Moment of inertia (mm<sup>4</sup>) | Section modulus (mm<sup>3</sup>) | Whole bone strength (N*mm) | Distance from neutral axis (mm) | Femur length (mm) | Tissue strength (N/mm<sup>2</sup>) |
| --- | --- | --- | --- | --- | --- | --- | --- | --- | --- |
| Body mass (g) |  | <u>**0.57**</u> | <u>**0.65**</u> | <u>**0.73**</u> | <u>**0.76**</u> | <u>**0.46**</u> | 0.29 | 0.04 | 0.04 |
| Fat pad mass (g) | [0.28, 0.77]<br>p < 0.001 |  | <u>**0.42**</u> | <u>**0.43**</u> | <u>**0.41**</u> | 0.24 | 0.22 | -0.04 | 0.03 |
| Cross-sectional area (mm<sup>2</sup>) | [0.39, 0.82]<br>p < 0.001 | [0.08, 0.67]<br>p = 0.018 |  | <u>**0.75**</u> | <u>**0.78**</u> | <u>**0.57**</u> | 0.31 | -0.02 | 0.14 |
| Moment of inertia (mm<sup>4</sup>) | [0.52, 0.86]<br>p < 0.001 | [0.09, 0.67]<br>p = 0.015 | [0.54, 0.87]<br>p < 0.001 |  | <u>**0.95**</u> | <u>**0.49**</u> | <u>**0.63**</u> | 0.28 | -0.06 |
| Section modulus (mm<sup>3</sup>) | [0.57, 0.88]<br>p < 0.001 | [0.07, 0.67]<br>p = 0.019 | [0.60, 0.89]<br>p < 0.001 | [0.90, 0.98]<br>p < 0.001 |  | <u>**0.44**</u> | <u>**0.36**</u> | 0.20 | -0.14 |
| Whole bone strength (N*mm) | [0.13, 0.70]<br>p = 0.008 | [-0.12, 0.54]<br>p = 0.190 | [0.28, 0.77]<br>p < 0.001 | [0.17, 0.72]<br>p = 0.004 | [0.11, 0.68]<br>p = 0.012 |  | 0.35 | -0.05 | <u>**0.82**</u> |
| Distance from neutral axis (mm) | [-0.06, 0.58]<br>p = 0.100 | [-0.13, 0.53]<br>p = 0.220 | [-0.05, 0.59]<br>p = 0.086 | [0.36, 0.80]<br>p < 0.001 | [0.02, 0.63]<br>p = 0.040 | [-0.001, 0.62]<br>p = 0.051 |  | 0.32 | 0.14 |
| Femur length (mm) | [-0.32, 0.40]<br>p = 0.830 | [-0.39, 0.33]<br>p = 0.850 | [-0.37, 0.35]<br>p = 0.930 | [-0.9, 0.58]<br>p = 0.130 | [-0.17, 0.52]<br>p = 0.290 | [-0.40, 0.31]<br>p = 0.790 | [-0.04, 0.61]<br>p = 0.081 |  | -0.23 |
| Tissue strength (N/mm<sup>2</sup>) | [-0.31, 0.38]<br>p = 0.830 | [-0.32, 0.37]<br>p = 0.890 | [-0.22, 0.46]<br>p = 0.450 | [-0.40, 0.29]<br>p = 0.720 | [-0.47, 0.21]<br>p = 0.430 | [0.66, 0.91]<br>p < 0.001 | [-0.22, 0.46]<br>p = 0.450 | [-0.54, 0.15]<br>p = 0.230 |  |

**Supplementary Table 5:** Pearson correlations between body mass, fat pad mass, biomechanics, and femoral geometry in aged female mice. The upper right shows r values while the bottom left shows confidence intervals (95%) and p-values are shown (bold and underlined: p < 0.05).

|  | Body mass<br>(g) | Fat pad<br>mass<br>(g) | Cross-sectional<br>area<br>(mm<sup>2</sup>) | Moment<br>of<br>inertia<br>(mm<sup>4</sup>) | Section<br>modulus<br>(mm<sup>3</sup>) | Whole<br>bone<br>strength<br>(N*mm) | Distance<br>from<br>neutral<br>axis<br>(mm) | Femur<br>length<br>(mm) | Tissue<br>strength<br>(N/mm<sup>2</sup>) |
| --- | --- | --- | --- | --- | --- | --- | --- | --- | --- |
| Body mass<br>(g) |  | **<u>0.87</u>** | **<u>0.67</u>** | **<u>0.71</u>** | **<u>0.65</u>** | 0.37 | **<u>0.60</u>** | 0.18 | 0.11 |
| Fat pad<br>mass<br>(g) | [0.69, 0.95]<br>$p < 0.001$ | | **<u>0.58</u>** | **<u>0.58</u>** | **<u>0.59</u>** | 0.33 | 0.41 | 0.05 | 0.11 |
| Cross-sectional<br>area<br>(mm<sup>2</sup>) | [0.33, 0.86]<br>$p = 0.001$ | [0.18, 0.81]<br>$p = 0.008$ | | **<u>0.87</u>** | **<u>0.81</u>** | **<u>0.70</u>** | **<u>0.70</u>** | 0.31 | 0.42 |
| Moment of<br>inertia<br>(mm<sup>4</sup>) | [0.39, 0.88]<br>$p < 0.001$ | [0.18, 0.81]<br>$p = 0.008$ | [0.69, 0.95]<br>$p < 0.001$ | | **<u>0.92</u>** | **<u>0.47</u>** | **<u>0.86</u>** | 0.33 | 0.11 |
| Section<br>modulus<br>(mm<sup>3</sup>) | [0.30, 0.85]<br>$p = 0.002$ | [0.20, 0.82]<br>$p = 0.006$ | [0.58, 0.92]<br>$p < 0.001$ | [0.81, 0.97]<br>$p < 0.001$ | | **<u>0.49</u>** | **<u>0.60</u>** | 0.30 | 0.10 |
| Whole<br>bone<br>strength<br>(N*mm) | [-0.09, 0.70]<br>$p = 0.110$ | [-0.13, 0.67]<br>$p = 0.150$ | [0.38, 0.87]<br>$p < 0.001$ | [0.04, 0.76]<br>$p = 0.036$ | [0.06, 0.77]<br>$p = 0.028$ | | 0.32 | 0.18 | **<u>0.91</u>** |
| Distance<br>from<br>neutral<br>axis<br>(mm) | [0.21, 0.82]<br>$p = 0.005$ | [-0.04, 0.72]<br>$p = 0.073$ | [0.38, 0.87]<br>$p < 0.001$ | [0.67, 0.94]<br>$p < 0.001$ | [0.22, 0.82]<br>$p = 0.005$ | [-0.15, 0.66]<br>$p = 0.170$ | | 0.29 | 0.07 |
| Femur<br>length<br>(mm) | [-0.28, 0.58]<br>$p = 0.440$ | [-0.40, 0.48]<br>$p = 0.840$ | [-0.15, 0.66]<br>$p = 0.180$ | [-0.13, 0.67]<br>$p = 0.160$ | [-0.16, 0.66]<br>$p = 0.190$ | [-0.29, 0.57]<br>$p = 0.450$ | [-0.17, 0.65]<br>$p = 0.210$ | | 0.04 |
| Tissue<br>strength<br>(N/mm<sup>2</sup>) | [-0.35, 0.53]<br>$p = 0.630$ | [-0.35, 0.53]<br>$p = 0.630$ | [-0.03, 0.73]<br>$p = 0.066$ | [-0.35, 0.52]<br>$p = 0.650$ | [-0.36, 0.52]<br>$p = 0.680$ | [0.78, 0.96]<br>$p < 0.001$ | [-0.38, 0.50]<br>$p = 0.770$ | [-0.41, 0.47]<br>$p = 0.870$ | |

**Supplementary Table 6:**
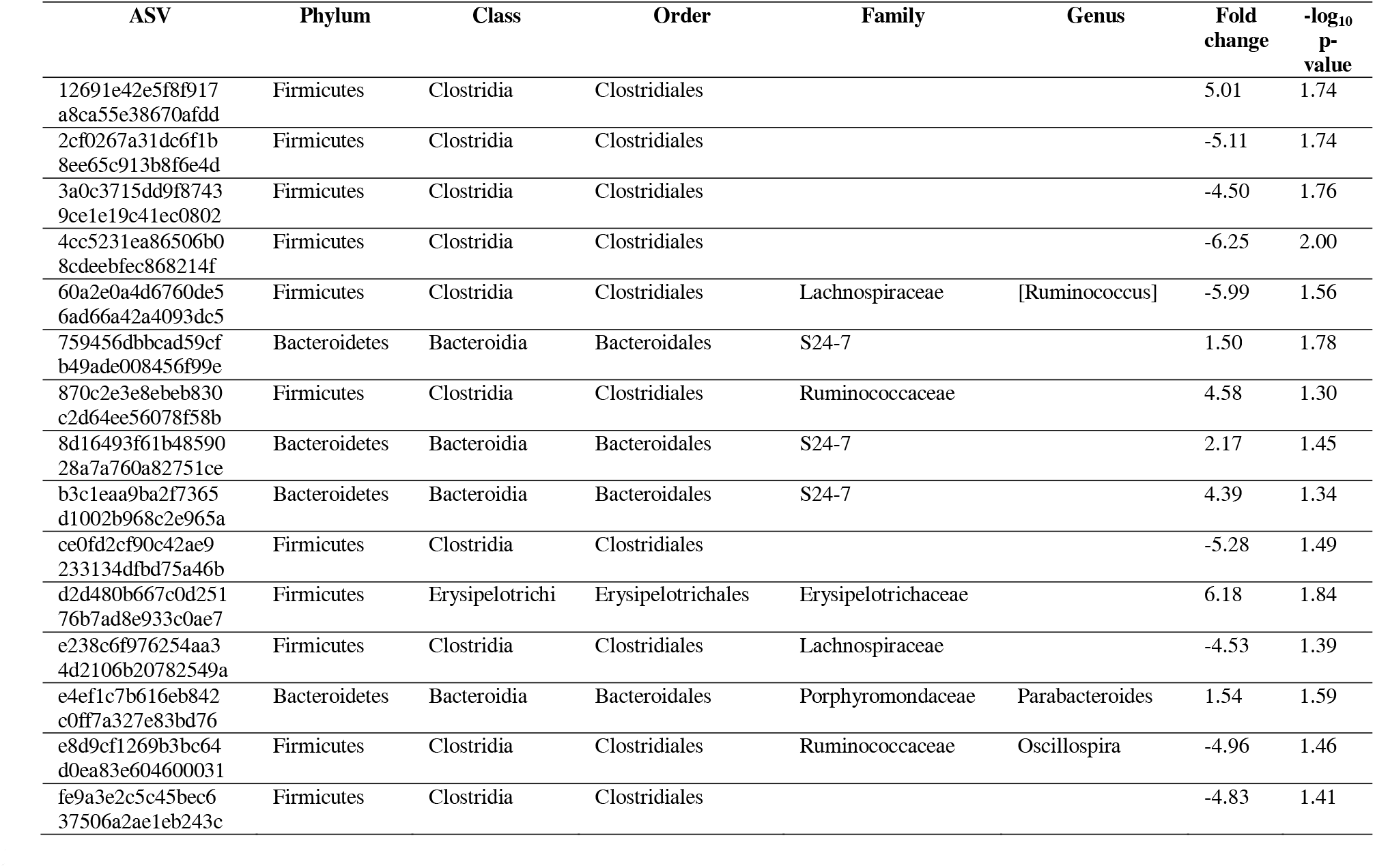
Taxonomy of significant ASVs (p < 0.05, magnitude of fold change >|1|) in aged males identified using ALDEx2.

**Supplementary Table 7:**
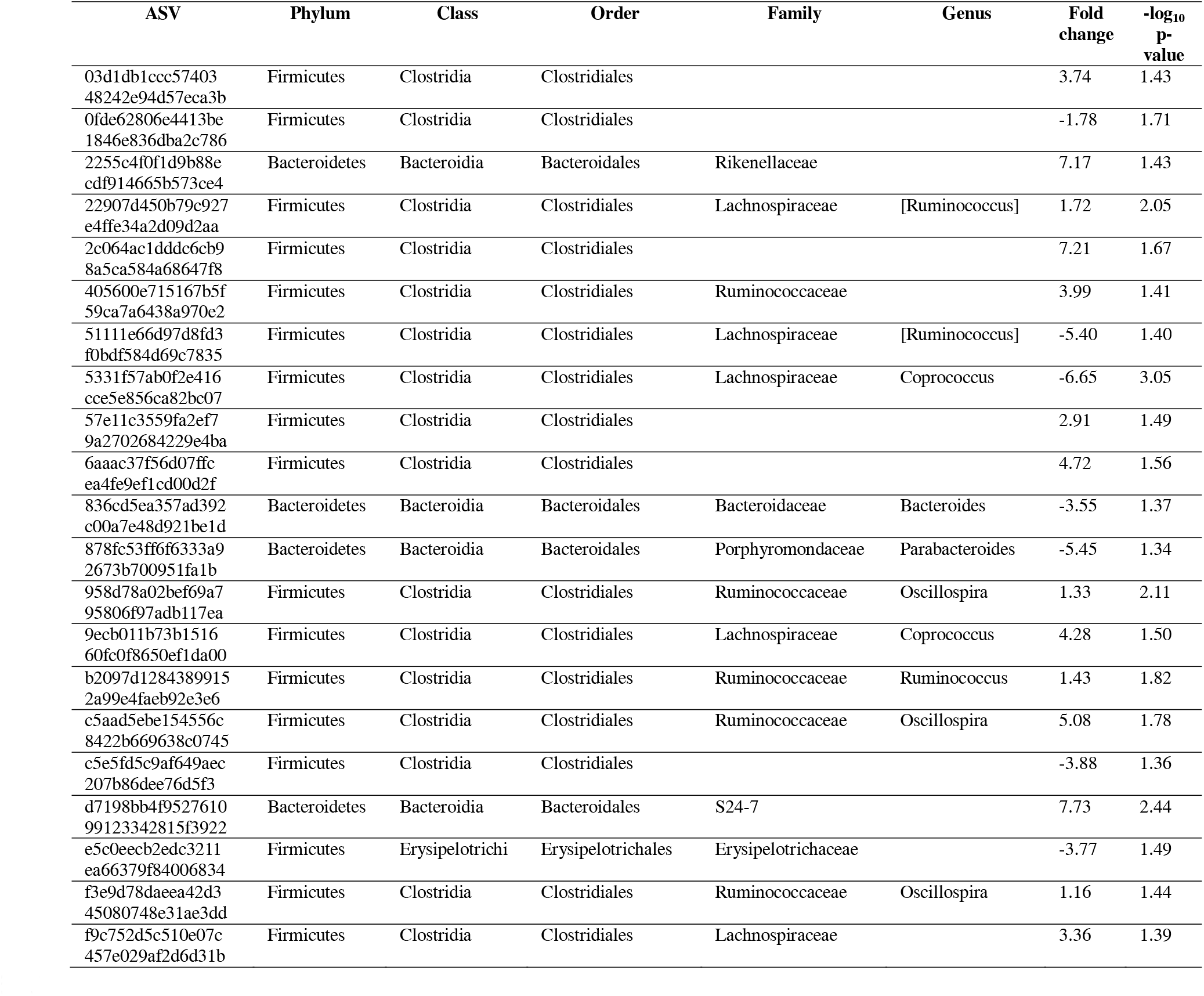
Taxonomy of significant ASVs (p < 0.05, magnitude of fold change > |1|) in aged females identified using ALDEx2.

